## Extended view text and figures for "High-throughput phenotypic screen for genetic modifiers in patient-derived *OPA1* mutant fibroblasts identifies *PGS1* as a functional suppressor of mitochondrial fragmentation"

**Extended View**

**figure EV1: Mitochondrial morphology supervised machine learning pipeline**

**(a)** Mitochondrial morphology quantification workflow using PhenoLOGIC Harmony 4.9 supervised machine learning (Table S1). Fluorescence imaging of cells (1) followed by nuclei (2) and cell segmentation (3) using cytoplasm defined by mitochondria. Cells on the border of the field of view are excluded from downstream analyses (4). Mitochondrial signal intensity is transformed into various texture parameters (5). (6) Supervised ML is trained using cells harbouring mitochondria that are fragmented (*Opa1* siRNA, red), hypertubular (*Dnm1l* siRNA, blue) or normal (NT siRNA, green). Debris cells (yellow) not included in the classification. (7) Class segregation represented by goodness of fit and then applied to unknown, non-trained images (8).

**(b)** Mitochondrial morphology quantification of control (CTL-1) fibroblasts treated with indicated siRNAs at the indicated doses for 72 hours. Supervised ML training performed on cells with fragmented (*OPA1* siRNA), normal (non-targeting NT siRNA), and hypertubular (*DNM1L* siRNA) mitochondria. Data represent mean ± SD of 3 independent experiments, (1005 to 2803 cells per cell line per replicate), One-way ANOVA.

**(c)** Supervised ML performed on mitochondrial morphologies generated by chemical manipulation (fragmented, CCCP, hypertubular, cycloheximide (CHX)) classify Opa1 deficient MEFs ($Opa1^{Crispr}$) before and after stable functional complementation with of Opa1s1-Myc (pTW264) or Opa1s7-Myc (pTW265). Data represent mean ± SD of 3-6 independent wells (1693 to 2554 cells per cell line), One-way ANOVA (% normal).

**(d)** Equal amounts of protein extracted from control and (CTL1, 2, 3) and DOA+ patient fibroblasts were separated by SDS-PAGE, immunoblotted with anti-OPA1 antibody, and quantified by densitometry relative to Ponceau stain. Data represent mean ± SD of three independent experiments, One-way ANOVA (relative to mean of CTL1, 2 and 3).

**figure EV2: High throughput screening identifies known and novel genetic modifiers of mitochondrial morphology in control and DOA+ patient-derived fibroblasts**

Representative confocal images of candidate genes (siRNAs) able to fragment **(a)** or hypertubulate **(b)** mitochondrial morphology identified in *Mitome* screen of *control* (CTL-1) fibroblasts (Figure 2A-C)

**(c)** String analysis clustering of 22 candidate genes that stimulate mitochondrial fragmentation upon depletion (Table S2)

**(d)** String analysis clustering of 145 candidate genes that stimulate mitochondrial hypertubulation upon depletion (Table S2), including clusters related to mitochondrial dynamics (top inset) and protein translation (bottom inset), (Table S2).

Representative confocal images of candidate genes (siRNAs) able to rescue **(e)** or hyperfragment **(f)** mitochondrial morphology identified in *Mitome* screen of $OPA1^{S545R}$ patient fibroblasts (Figure 2E-F).

**(g)** Violin plot representing % Hyperfragmented morphology of ground truth and *Mitome* siRNAs. The siRNA able to hyperfragment mitochondrial morphology in $OPA1^{S545R}$ patient fibroblasts were selected with a univariate 3-components statistical model programmed in R using ground truths for morphology in Figure 2E. The defined threshold for positive hits was 72.9% and identified 27 candidate genes (Table S3).

Functional classification **(h)** and String analysis clustering **(i)** of 91 candidate genes that rescued mitochondrial fragmentation in $OPA1^{S545R}$ fibroblasts upon depletion (Table S3).

**figure EV3: PGS1 depletion rescues mitochondrial fragmentation in OPA1-deficient human and mouse fibroblasts**

**(a)** *Opa1* splice isoforms in MEFs. Proteolytic cleavage sites for MPP, OMA1, and YME1L are indicated. TM, transmembrane domain.

**(b)** Equal amounts of protein extracted from WT, *Opa1*$^{Crispr}$ and *Opa1*$^{KO}$ MEFs were separated by SDS-PAGE, immunoblotted with indicated antibodies.

**(c)** quantitative RT-PCR (qRT-PCR) measurement of *Opa1* and *Pgs1* expression in indicated *Opa1*$^{Crispr}$ MEFs complemented with pLenti-Opa1 and *Opa1*$^{Crispr}$*Pgs1*$^{Crispr}$ MEFs complemented with pLenti-Pgs1 relative to WT control. Data represent mean ± SD of two independent experiments, unpaired t-test.

**(d)** Representative confocal images of WT, *Opa1*$^{Crispr}$, and *Opa1*$^{Crispr}$ MEFs complemented with pLenti-Opa1-Myc. Mitochondria (anti-TOM40, green) and nuclei (DAPI, blue) and Opa1-Myc (anti-Myc, red). Scale bar=10μm

**(e)** Mitochondrial morphology quantification of (d) using WT MEFs treated with *Opa1* siRNA (fragmented), NT siRNA (normal), or *Dnm1l* siRNA (hypertubular) ground truth training sets. Data represent mean ± SD of six independent experiments, (1693-3200 cells per cell line), One-way ANOVA (% fragmented).

**(f)** Equal amounts of protein extracted from MEFs in (E) were separated by SDS-PAGE, immunoblotted with indicated antibodies.

**(g)** Mitochondrial membrane potential measured in WT, *Opa1*$^{Crispr}$, *Opa1*$^{Crispr}$ + pLenti-Opa1-Myc MEFs abeled with TMRE and analyzed by cytometry. TMRE/mitoYFP was used to normalize membrane potential. Data represent mean ± SD of three independent experiments (>10,000 cells/sample), One-way ANOVA.

**(h)** Representative confocal images of wild type (WT) and *Opa1*$^{Crispr}$ MEFs treated with NT or *Opa1* siRNA for 72 hours. Live imaging of mitochondria (mitoYFP, green) and nuclei (NucBlue, blue). Scale bar=10μm.

**(i)** Mitochondrial morphology quantification of (H) using WT MEFs treated with *Opa1* siRNA (fragmented), NT siRNA (normal), or *Dnm1l* siRNA (hypertubular) training sets. Data represent mean ± SD of three independent experiments, (550-822 cells per cell line), One-way ANOVA (% fragmented).

**(j)** Representative confocal images of wild type (WT) and *Opa1*$^{Crispr}$ MEFs treated with 50 µM cycloheximide (CHX) for 3 hours. Live imaging of mitochondria (mitoYFP, green) and nuclei (NucBlue, blue). Scale bar=10µm.

**(k)** Mitochondrial morphology quantification of (J) using WT MEFs treated with *Opa1* siRNA (fragmented), NT siRNA (normal), or *Dnm1l* siRNA (hypertubular) training sets. Data represent mean ± SD of three independent experiments, (547-822 cells per cell line), One-way ANOVA (% fragmented).

**(l)** Representative confocal images of wild type (WT) and *Opa1*$^{KO}$ MEFs treated with 50 µM cycloheximide (CHX) for 3 hours. Live imaging of mitochondria (mitoYFP, green) and nuclei (NucBlue, blue). Scale bar=10µm.

**(m)** Mitochondrial morphology quantification of (K) using WT MEFs treated with *Opa1* siRNA (fragmented), NT siRNA (normal), or *Dnm1l* siRNA (hypertubular) training sets. Data represent mean ± SD of two independent experiments, (638-1063 cells per cell line) One-way ANOVA (% hypertubular).

**figure EV4: PGS1 depletion rescues mitochondrial fragmentation by inhibiting mitochondrial fission independently of Drp1 recruitment and Opa1 processing.**

**(a)** Equal amounts of protein extracted from WT and mutant MEFs stably expressing pLenti-Opa1 or pLenti-Pgs1 where indicated were separated by SDS-PAGE, immunoblotted with indicated antibodies and quantified by densitometry **(b)** relative to tubulin. Data represent mean ± SD of four independent experiments,

**(c)**Equal amounts of protein extracted from WT MEFs treated with 20 µM CCCP (30min) or 16 µM 4Br-A23187 (18h) were separated by SDS-PAGE, immunoblotted with indicated antibodies. Tubulin was used as loading control.

**(d)** Representative confocal images of live cell imaging of MEFs of the indicated genotypes subjected to fission with 4Br-A23187 for the indicated time points. Images were captured every hour for 18 hours. Scale bar=10μm.

**(e)** Mitochondrial morphology quantification of (d) using WT MEFs treated with 5 μM CCCP for 18h (fragmented), untreated (normal), or treated with 10 μM CHX for 9h (hypertubular) training sets. Data represent mean ± SD of 3 independent experiments, (355-1196 cells per cell line) One-way ANOVA.

**(f)** Quantification of mitochondrial membrane potential (TMRE/mitoYFP) by confocal microscopy in WT and *Pgs1*$^{Crispr}$ MEFs treated as indicated. Number of analyzed cells indicated within bar. Data represent mean ± SEM of four independent experiments, One-way ANOVA.

**(g)** Equal amounts of protein extracted from WT and mutant MEFs treated with CCCP at the indicated concentrations and durations where indicated were separated by SDS-PAGE, immunoblotted with indicated antibodies and quantified by densitometry.

**(h)** Representative confocal images of live cell imaging of MEFs of the indicated genotypes subjected to stress-induced mitochondrial hyperfusion (SiMH) with 0.5 μM ActD for the indicated time points. Images were captured every 20 minutes for 240 minutes. Scale bar=10μm

**(I)** Mitochondrial morphology quantification of using WT MEFs treated with 5 μM CCCP for 18h (fragmented), untreated (normal), or treated with 10 μM CHX for 9h (hypertubular) training sets. Data represent mean ± SD of 3 independent experiments, (262-1123 cells per cell line) One-way ANOVA.

**figure EV5: Cardiolipin remodelling in Opa1 and Pgs1-deficient fibroblasts**

**(a)** Representative confocal micrographs of MEFs WT and *Opa1*$^{KO}$ MEFs treated with indicated siRNAs for 72 hours. Mitochondria (anti-TOM40, green) and nuclei (DAPI, blue). Scale bar=10μm.

**(b)** Mitochondrial morphology quantification of (a) using WT MEFs treated with *Opa1* siRNA (fragmented), NT siRNA (normal), or *Dnm1l* siRNA (hypertubular) training sets. Data represent mean ± SD of two independent experiments, (185-2689 cells per cell line), One-way ANOVA (% fragmented).

**(c)** Cardiolipin (CL) saturation state (carbon double bonds per DAG molecule) determined from purified mitochondria isolated from MEFs of the indicated genotypes. Data represent mean ± SD of four independent experiments, One-way ANOVA.

**(d)** CL acyl chain composition determined from purified mitochondria isolated from MEFs of the indicated genotypes. Data represent mean ± SD of four independent experiments.

**figure EV6: Pgs1 depletion does not restore apoptotic phenotype of Opa1-deficient MEFs.**

**(a, b)** MEFs of the indicated genotypes were subjected to 0.5 μM staurosporine in the presence or absence of the pan-caspase inhibitor qVD. Dead cells (PI+ nuclei, orange) and total cells (NucBlue, blue) were imaged every hour for 25 hours. PI+ nuclei number divided by the total nuclei number was then quantified over time. (Bottom) Representative confocal images of (Top). Scale bar=100μm. Data represent mean ± SD of four independent experiments, 1923-4703 cells per cell line) One-way ANOVA.

**(c, d)** MEFs of the indicated genotypes were subjected to 16 μM etoposide. Dead cells (PI+ nuclei, orange) and total cells (NucBlue, blue) were imaged every hour for 25 hours. PI+ nuclei number divided by the total nuclei number was then quantified over time. (Bottom)

Representative confocal images of (Top). Scale bar=100µm. Data represent mean ± SD of four independent experiments, 1816-3915 cells per cell line) One-way ANOVA.

**figure EV7: Opa1-Myc does not complement respiration defect in Opa1-deficient MEFs.**

(**a**) Basal oxygen consumption (O2 flux) measured by O2k high resolution respirometry (Oroboros) in WT, *Opa1*$^{Crispr}$, *Opa1*$^{Crispr}$ + pLenti-Opa1, *Opa1*$^{Crispr}$*Pgs1*$^{Crispr}$, *Opa1*$^{Crispr}$*Pgs1*$^{Crispr}$ + pLenti-Pgs1, *Pgs1*$^{Crispr}$ and *Pgs1*$^{Crispr}$ MEFs + pLenti-Pgs1. Measurements were made in pairwise fashion compared to WT MEFs. O2 flux normalized to protein concentration. Data represent mean ± SD of three independent experiments, unpaired t-test.

Mitochondrial respiration measured in adherent MEFs of the indicated genotypes using Seahorse FluxAnalyzer. Oxygen consumption rate (OCR) normalized to protein concentration. Following basal respiration, cells were treated sequentially with 1 µM Oligomycin (Omy), 2 µM CCCP (maximal). Bar graphs of representing basal **(b)** and maximal **(c)** respiration. Data represent mean ± SEM of 22-24 parallel OCR measurement, One-way ANOVA.

**Supplemental tables**

**Table S1:** Mitochondrial morphology quantification and Drp1 colocalization workflows on Harmony 4.9

**Table S2:** Mitome siRNA library contents and plate layouts

**Table S3:** Mitome siRNA candidate genes identified in CTL-1 and CTL-2 control human fibroblasts that lead to mitochondrial hypertubulation or fragmentation.

**Table S4:** Mitome siRNA candidate genes identified in *OPA1*$^{S545R}$ patient-derived fibroblasts that lead to mitochondrial morphology rescue.

**Table S5:** Reagents used in this study including siRNAs, primers, and plasmids

**Movies:**FRAP fusion assay in mitoYFP WT (1), $Opa1^{Crispr}$ (2) and $Opa1^{Crispr}Pgs1^{Crispr}$ (3) MEFs imaged in Figure 4E every 200ms. Movies represented at 5 frames per second (see Supplemental Movies 1-3).

**Computer Script:** OPA1S545R -OPA1.Rmd univariate script in R used to define Mitome hit thresholds in Figure 2.

Table 1

| Patients (gender, age) | Age of onset | Optic Atrophy | CPEO | Ataxia | Spasticity | Peripheral Neuropathy | Deafness | *OPA1* Mutation | *OPA1* Domain | Reference |
|---|---|---|---|---|---|---|---|---|---|---|
| **Patient 1** (M, 30 years) | Childhood | + | - | + | - | + | + | m.1635C>G p.S545R | Dynamin | Yu-Wai-Man *et al* 2010 (Patient FR-1) |
| **Patient 2** (F, 37 years) | 6 years | + | - | + | - | - | + | m.1334G>A p.R445H | GTPase | Amati-Bonneau *et al* 2005 (Patient 1) |
| **Patient CS** (F, 60 years) | 50 years | + | - | + | + | - | - | c.2356-1G>T | Dynamin | Yu-Wai-Man *et al* 2016 (Patient A) |
| **Patient DM** (M, 43 years) | Childhood | + | + | + | + | - | - | c.1294A>G p.I432V | GTPase | Yu-Wai-Man *et al* 2010 (Patient UK-12) |
| **Patient CC** (F, 48 years) | < 5 years | + | - | - | + | + | - | c.899C>T p.Q297X | GTPase | Yu-Wai-Man *et al* 2010 (Patient UK-5) |

**Table 1:** Clinical features of patient fibroblasts. CPEO: chronic progressive external ophthalmoplegia; M: male; F: female.

Amati-Bonneau P, Guichet A, Olichon A, Chevrollier A, Viala F, Miot S, Ayuso C, Odent S, Arrouet C, Verny C, Calmels MN, Simard G, Belenguer P, Wang J, Puel JL, Hamel C, Malthièry Y, Bonneau D, Lenaers G, Reynier P. *OPA1* R445H mutation in optic atrophy associated with sensorineural deafness. Ann Neurol. 2005;58:958-63.

Yu-Wai-Man P, Griffiths PG, Gorman GS, Lourenco CM, Wright AF, Auer-Grumbach M, Toscano A, Musumeci O, Valentino ML, Caporali L, Lamperti C, Tallaksen CM, Duffey P, Miller J, Whittaker RG, Baker MR, Jackson MJ, Clarke MP, Dhillon B, Czermin B, Stewart JD, Hudson G, Reynier P, Bonneau D, Marques W Jr, Lenaers G, McFarland R, Taylor RW, Turnbull DM, Votruba M, Zeviani M, Carelli V, Bindoff LA, Horvath R, Amati-Bonneau P, Chinnery PF. Multi-system neurological disease is common in patients with *OPA1* mutations. Brain. 2010;133:771-86.

Yu-Wai-Man P, Spyropoulos A, Duncan JH, Guagdano JV, Chinnery PF. A multiple sclerosis-like disorder in patients with *OPA1* mutations. Annals of Clinical and Translational Neurology. 2016;3:723-9.

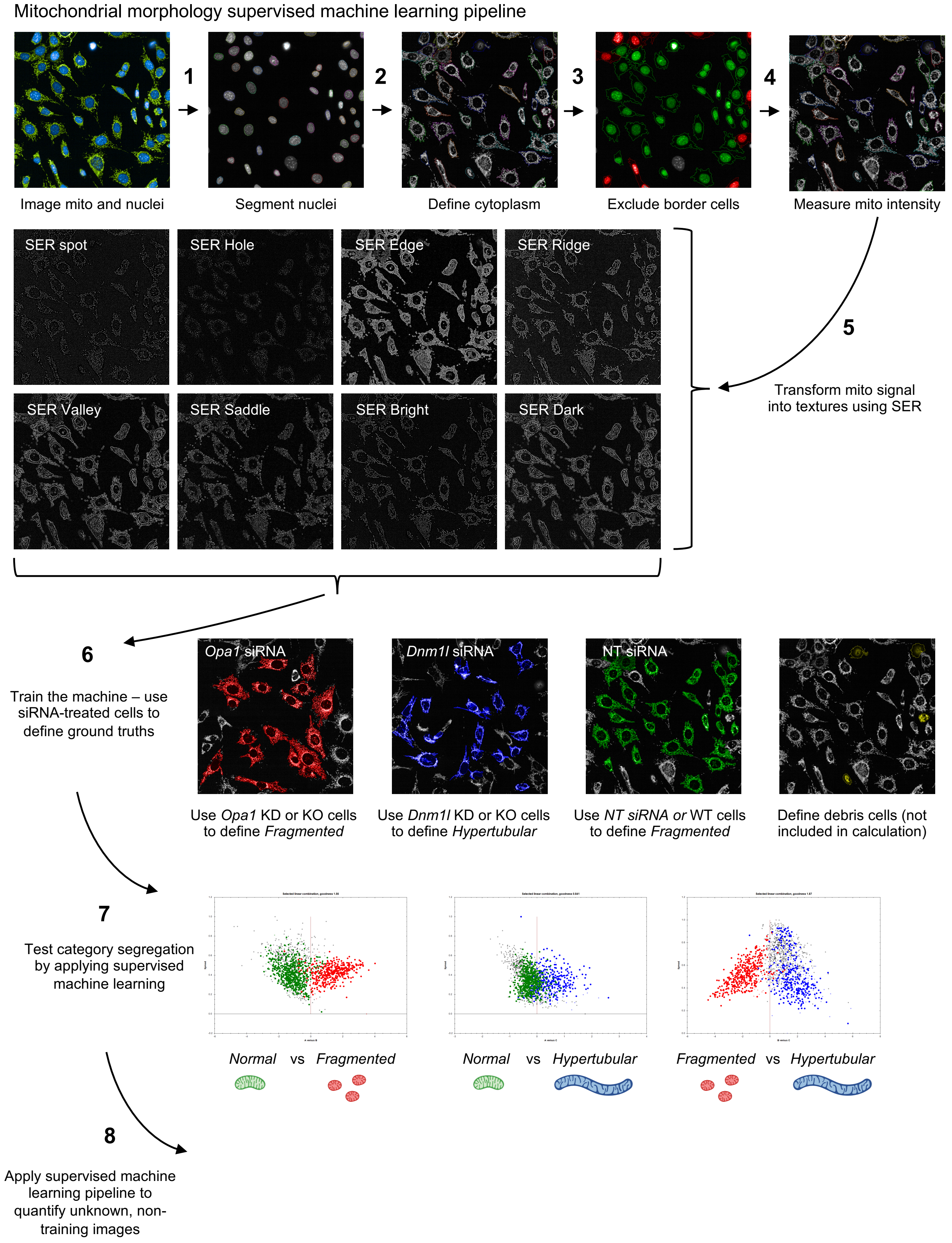
Mitochondrial morphology supervised machine learning pipeline
1
2
3
4
Image mito and nuclei
Segment nuclei
Define cytoplasm
Exclude border cells
Measure mito intensity
SER spot
SER Hole
SER Edge
SER Ridge
SER Valley
SER Saddle
SER Bright
SER Dark
5
Transform mito signal into textures using SER
6
Train the machine – use siRNA-treated cells to define ground truths
*Opa1* siRNA
*Dnm1l* siRNA
NT siRNA
Use *Opa1* KD or KO cells to define *Fragmented*
Use *Dnm1l* KD or KO cells to define *Hypertubular*
Use *NT siRNA or* WT cells to define *Fragmented*
Define debris cells (not included in calculation)
7
Test category segregation by applying supervised machine learning
*Normal* vs *Fragmented*
*Normal* vs *Hypertubular*
*Fragmented* vs *Hypertubular*
8
Apply supervised machine learning pipeline to quantify unknown, non-training images

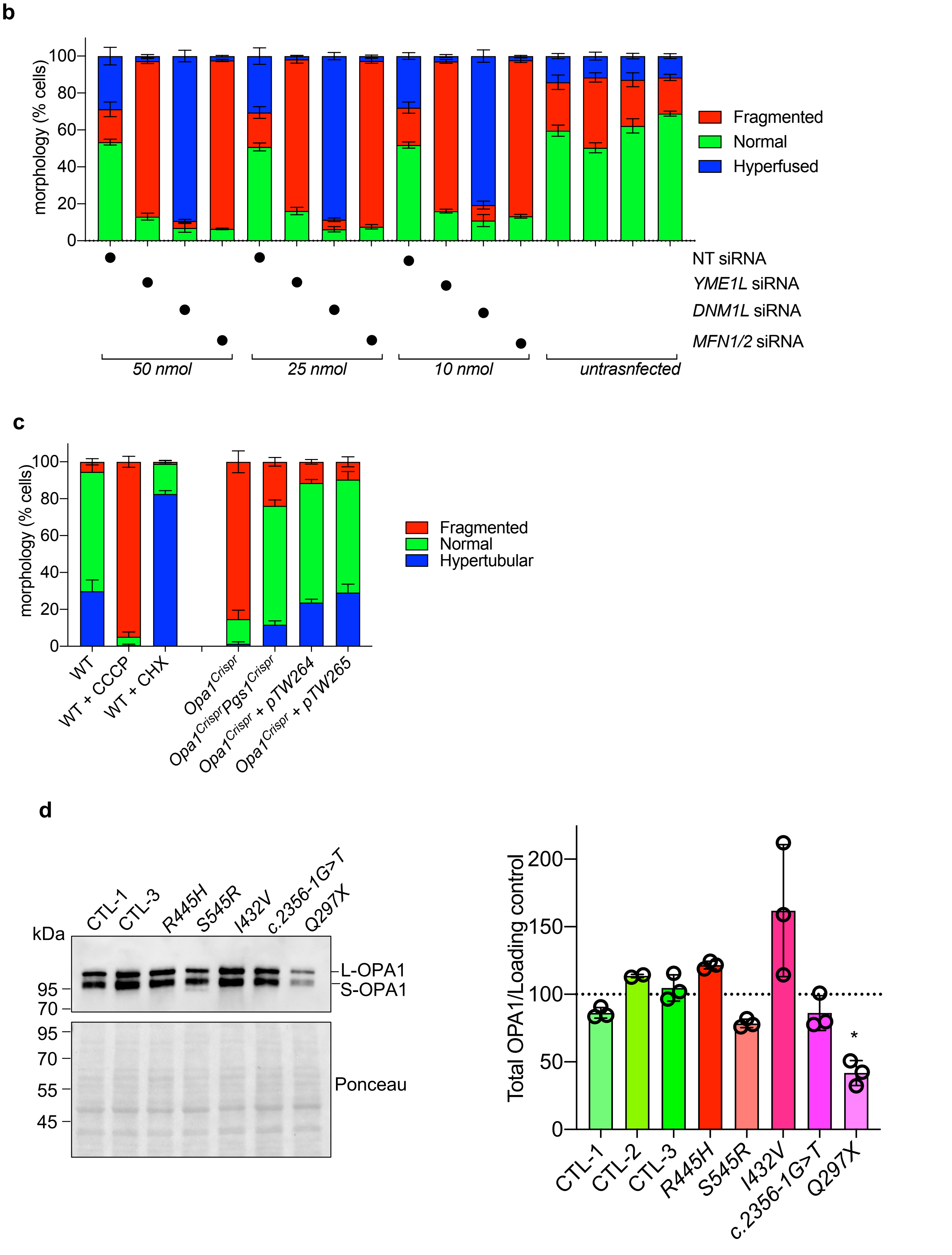
b
morphology (% cells)
0
20
40
60
80
100
Fragmented
Normal
Hyperfused
NT siRNA
YME1L siRNA
DNM1L siRNA
MFN1/2 siRNA
50 nmol
25 nmol
10 nmol
untrasnfected
c
morphology (% cells)
Fragmented
Normal
Hypertubular
WT
WT + CCCP
WT + CHX
Opa1Crispr
Opa1CrisprPgs1Crispr
Opa1Crispr + pTW264
Opa1Crispr + pTW265
d
kDa
CTL-1
CTL-3
R445H
S545R
I432V
c.2356-1G>T
Q297X
95
70
L-OPA1
S-OPA1
95
70
55
45
Ponceau
Total OPA1/Loading control
0
50
100
150
200
*
CTL-1
CTL-2
CTL-3
R445H
S545R
I432V
c.2356-1G>T
Q297X

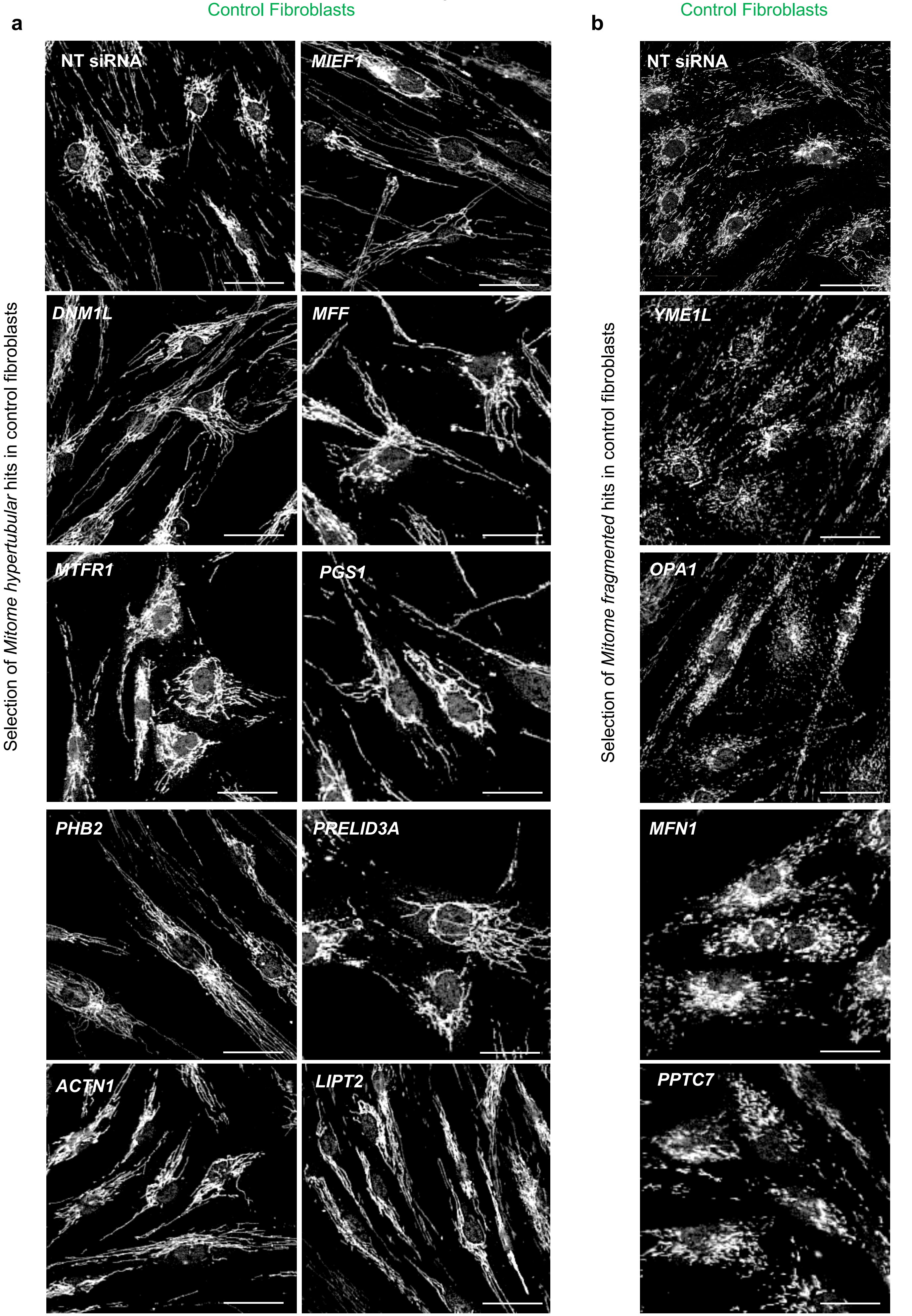
Control Fibroblasts
a
NT siRNA
MIEF1
DNM1L
MFF
MTFR1
PGS1
PHB2
PRELID3A
ACTN1
LIPT2
Selection of *Mitome hypertubular* hits in control fibroblasts
Control Fibroblasts
b
NT siRNA
YME1L
OPA1
MFN1
PPTC7
Selection of *Mitome fragmented* hits in control fibroblasts

**c** Mitochondrial fragmentation

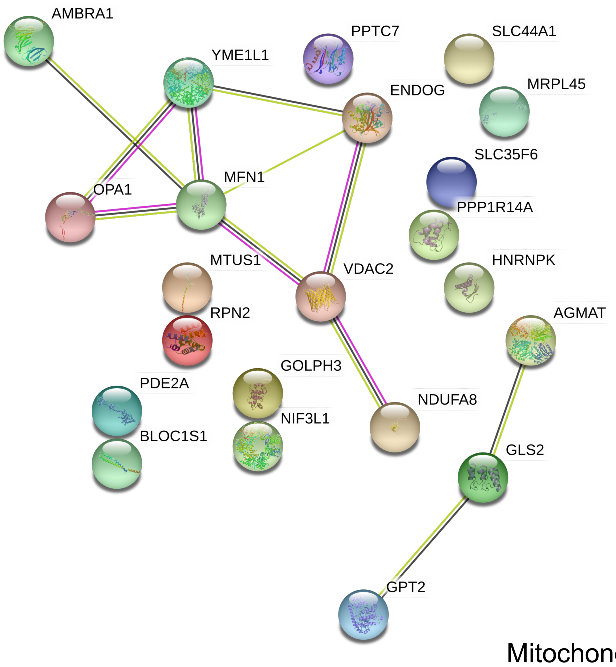


**d** Mitochondrial hypertubulation

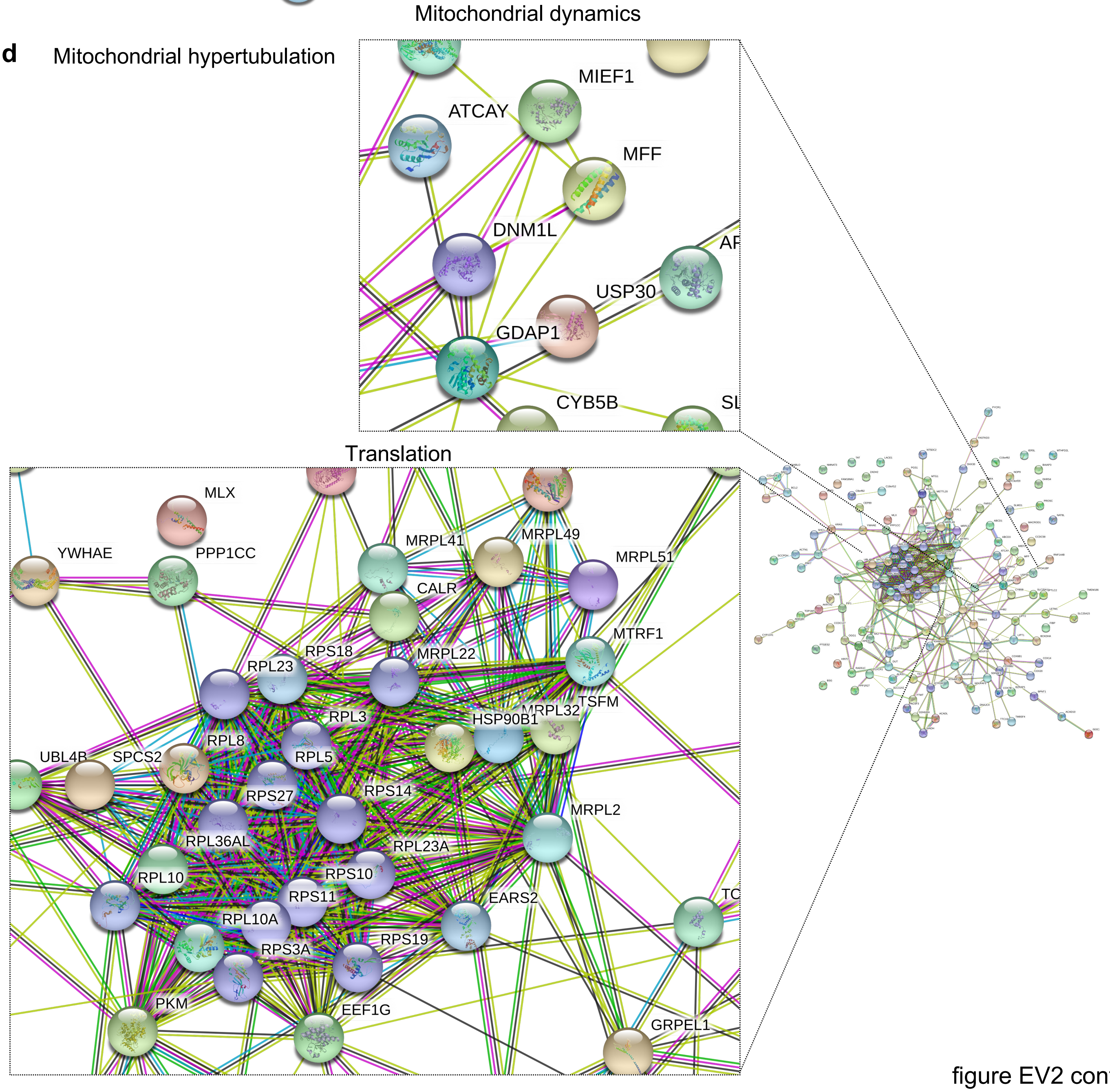

figure EV2 cont

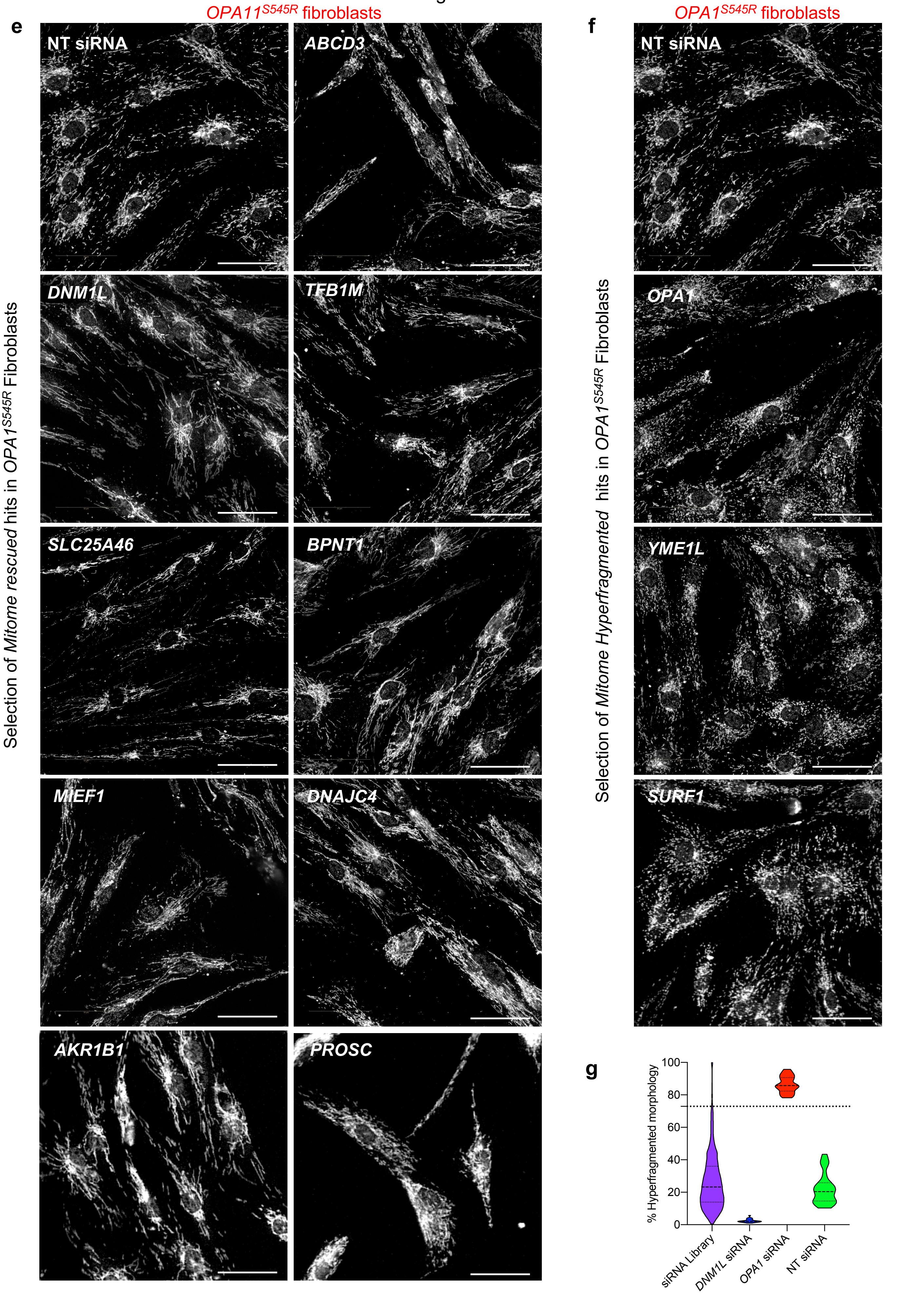


figure EV2 cont

h

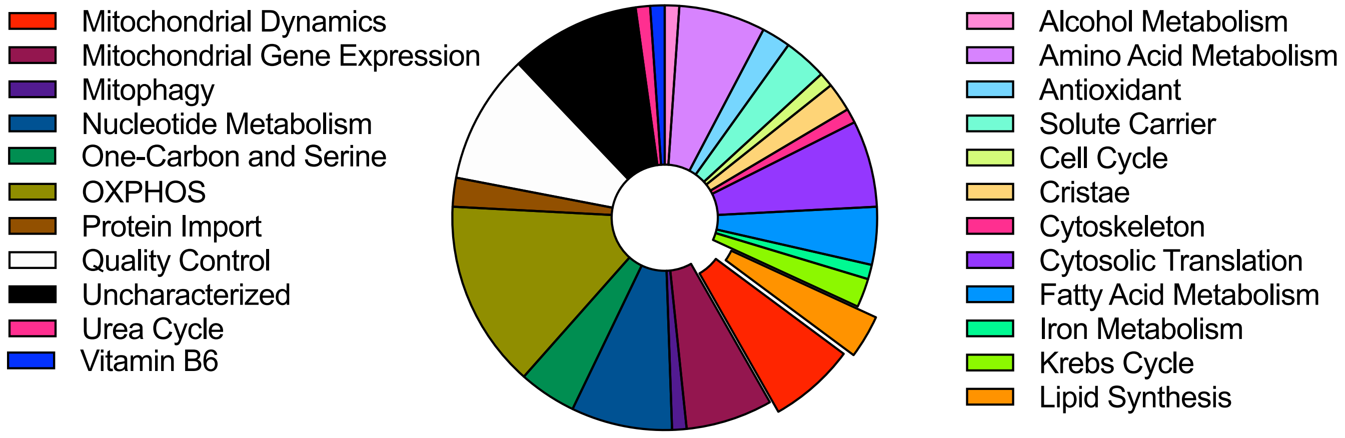
Mitochondrial Dynamics
Mitochondrial Gene Expression
Mitophagy
Nucleotide Metabolism
One-Carbon and Serine
OXPHOS
Protein Import
Quality Control
Uncharacterized
Urea Cycle
Vitamin B6
Alcohol Metabolism
Amino Acid Metabolism
Antioxidant
Solute Carrier
Cell Cycle
Cristae
Cytoskeleton
Cytosolic Translation
Fatty Acid Metabolism
Iron Metabolism
Krebs Cycle
Lipid Synthesis

i

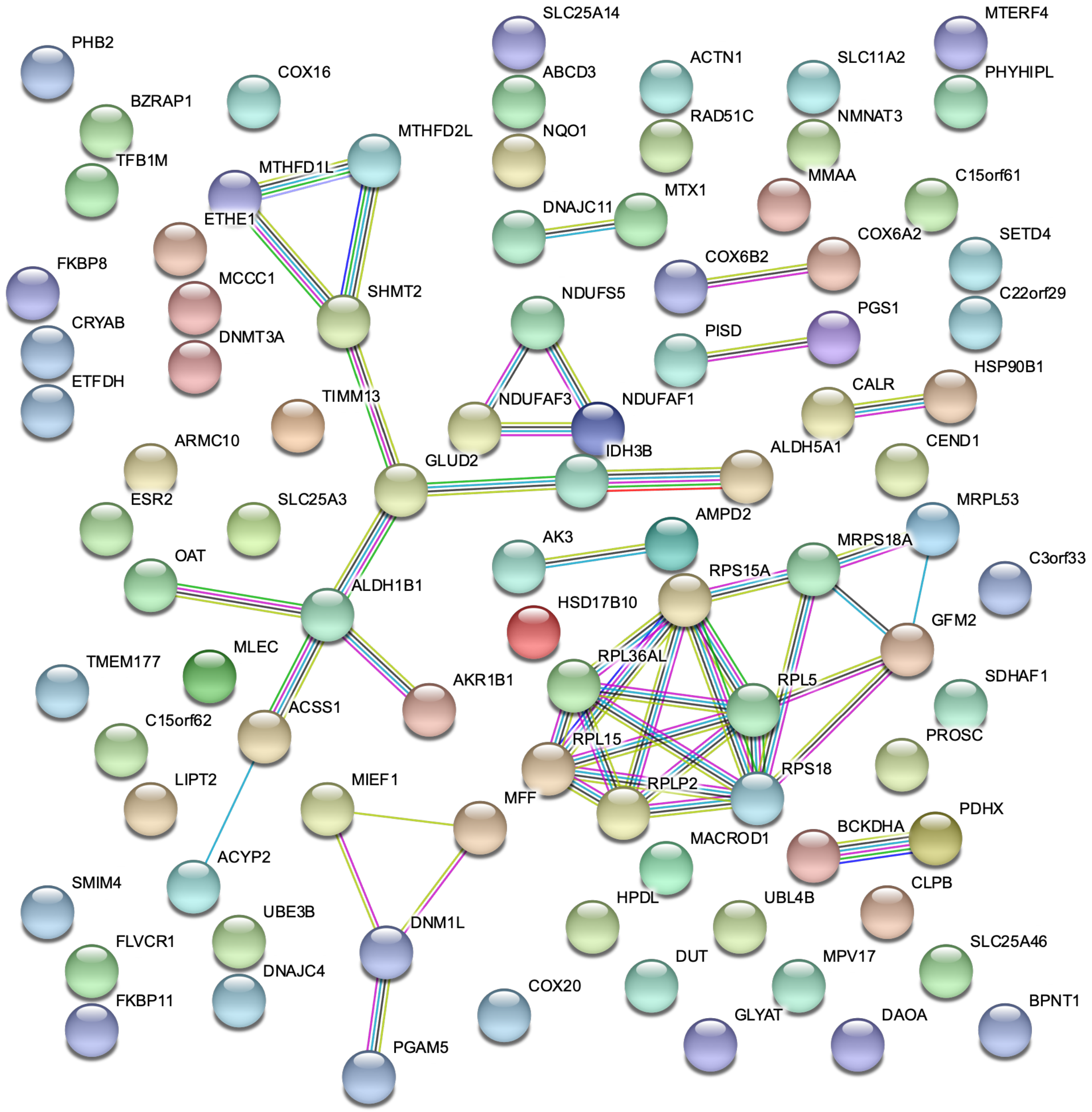
SLC25A14
MTERF4
PHB2
COX16
ABCD3
ACTN1
SLC11A2
PHYHIPL
BZRAP1
NQO1
RAD51C
NMNAT3
TFB1M
MTHFD2L
MTHFD1L
MMAA
C15orf61
MTX1
ETHE1
DNAJC11
COX6A2
SETD4
FKBP8
COX6B2
MCCC1
SHMT2
NDUFS5
C22orf29
CRYAB
PGS1
DNMT3A
PISD
ETFDH
HSP90B1
TIMM13
CALR
NDUFAF3
NDUFAF1
ARMC10
IDH3B
CEND1
GLUD2
ALDH5A1
ESR2
SLC25A3
MRPL53
AMPD2
OAT
AK3
MRPS18A
C3orf33
RPS15A
ALDH1B1
HSD17B10
GFM2
TMEM177
MLEC
RPL36AL
SDHAF1
AKR1B1
RPL5
ACSS1
C15orf62
PROSC
RPL15
LIPT2
MIEF1
RPS18
RPLP2
MFF
PDHX
MACROD1
BCKDHA
ACYP2
SMIM4
CLPB
HPDL
UBL4B
UBE3B
DNM1L
FLVCR1
SLC25A46
DUT
MPV17
DNAJC4
COX20
FKBP11
BPNT1
GLYAT
DAOA
PGAM5

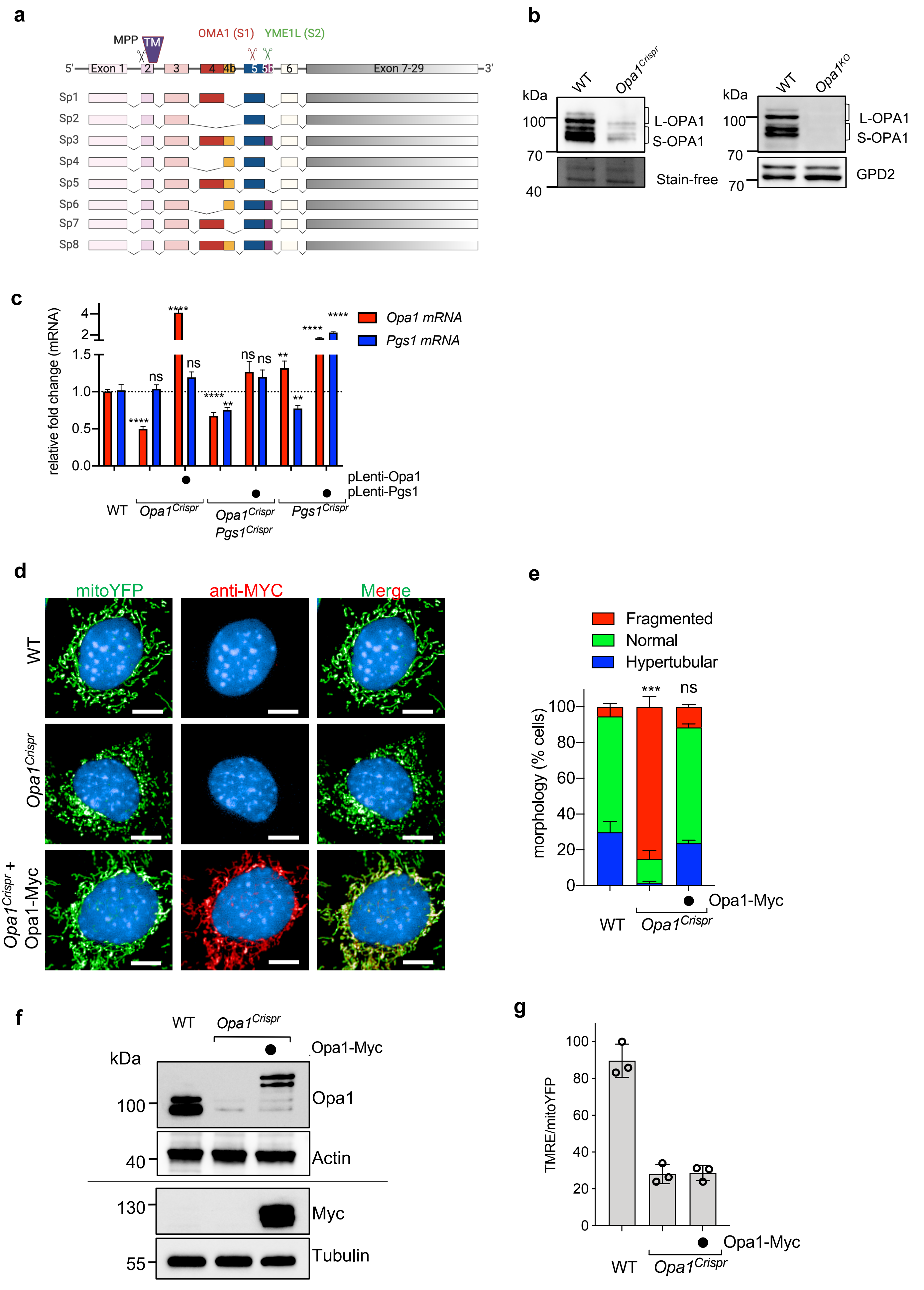
a
MPP
TM
OMA1 (S1)
YME1L (S2)
5'
Exon 1
2
3
4
4b
5
5b
6
Exon 7-29
3'
Sp1
Sp2
Sp3
Sp4
Sp5
Sp6
Sp7
Sp8
b
kDa
WT
Opa1Crispr
100
70
40
L-OPA1
S-OPA1
Stain-free
WT
Opa1KO
L-OPA1
S-OPA1
GPD2
c
relative fold change (mRNA)
Opa1 mRNA
Pgs1 mRNA
****
ns
**
pLenti-Opa1
pLenti-Pgs1
WT
Opa1Crispr
Opa1Crispr Pgs1Crispr
Pgs1Crispr
d
mitoYFP
anti-MYC
Merge
WT
Opa1Crispr
Opa1Crispr + Opa1-Myc
e
Fragmented
Normal
Hypertubular
***
ns
morphology (% cells)
Opa1-Myc
WT
Opa1Crispr
f
WT
Opa1Crispr
Opa1-Myc
kDa
100
40
130
55
Opa1
Actin
Myc
Tubulin
g
TMRE/mitoYFP
Opa1-Myc
WT
Opa1Crispr

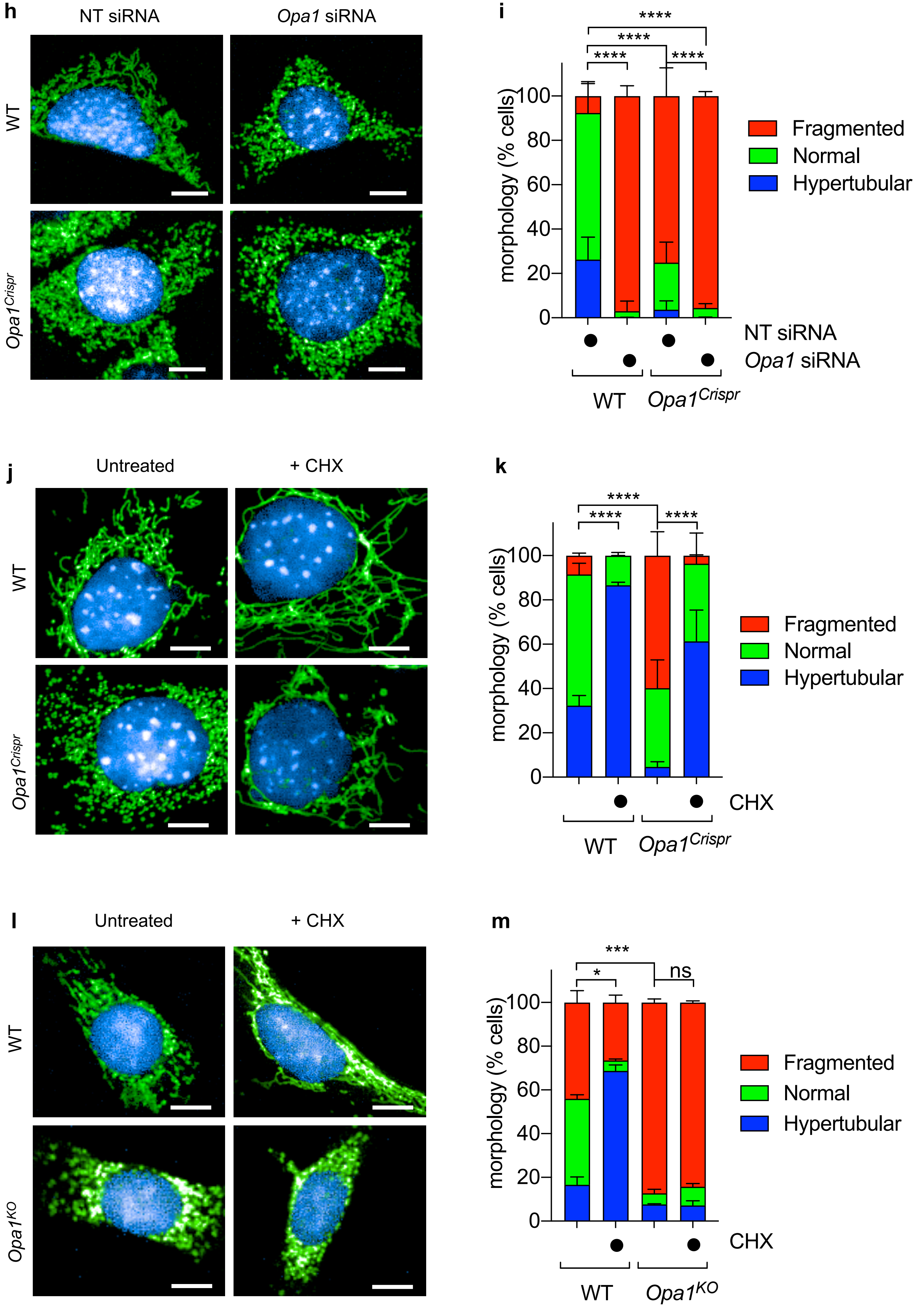
h
NT siRNA
*Opa1* siRNA
WT
*Opa1*Crispr
i
****
****
****
****
morphology (% cells)
100
80
60
40
20
0
Fragmented
Normal
Hypertubular
NT siRNA
*Opa1* siRNA
WT
*Opa1*Crispr
j
Untreated
+ CHX
WT
*Opa1*Crispr
k
****
****
****
morphology (% cells)
100
80
60
40
20
0
Fragmented
Normal
Hypertubular
CHX
WT
*Opa1*Crispr
l
Untreated
+ CHX
WT
*Opa1*KO
m
***
*
ns
morphology (% cells)
100
80
60
40
20
0
Fragmented
Normal
Hypertubular
CHX
WT
*Opa1*KO

figure EV4

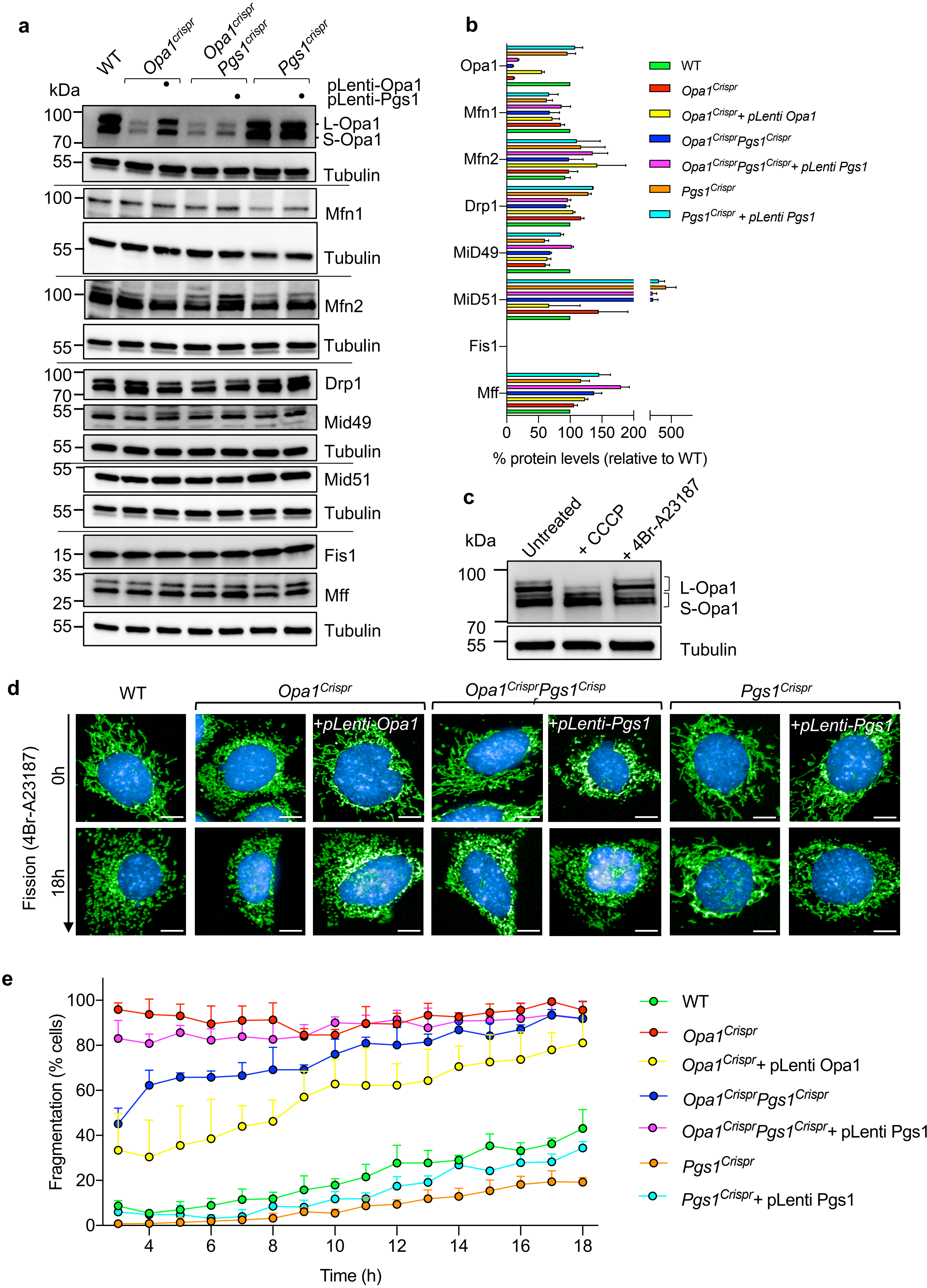


Figure EV4

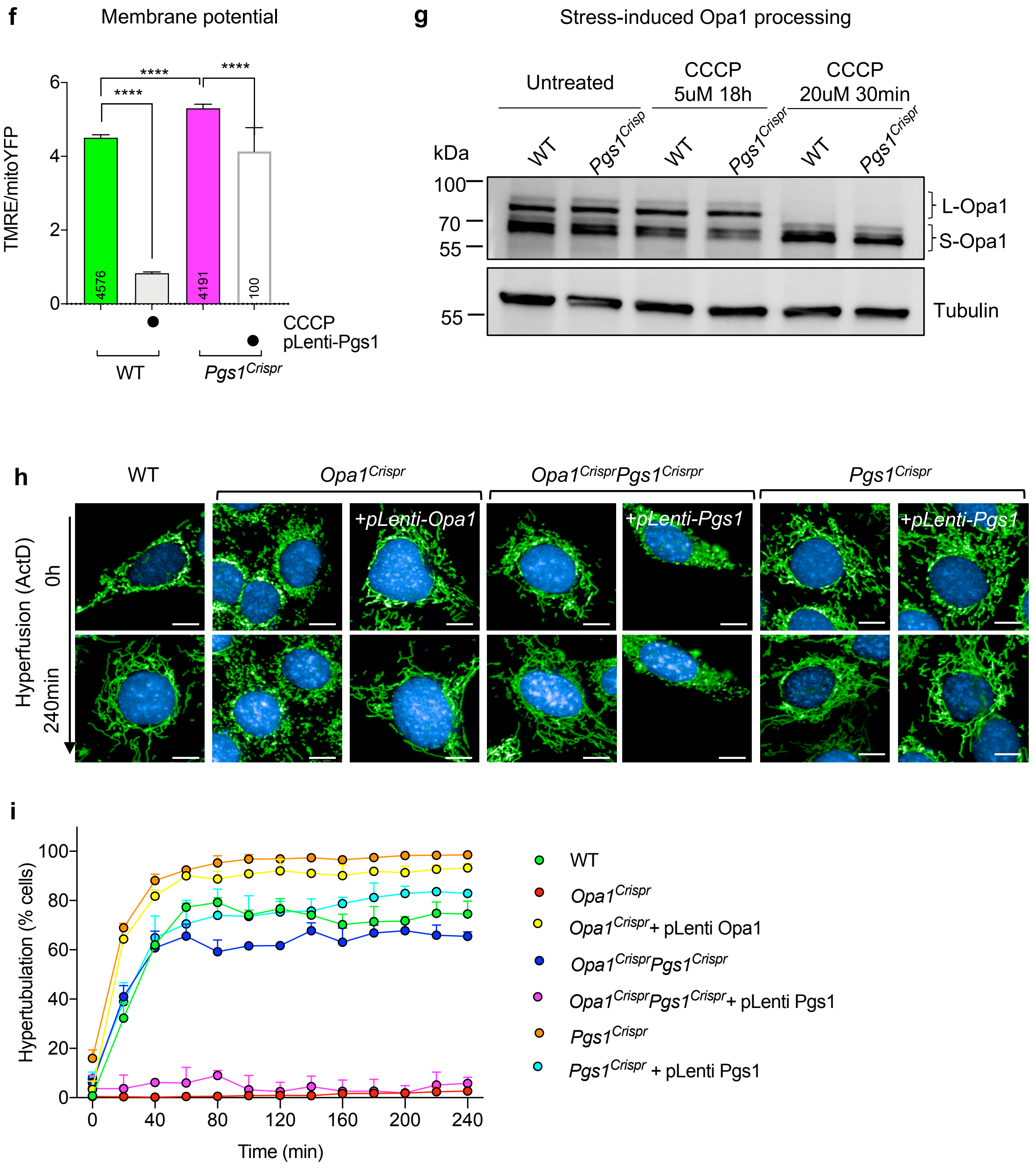
f
Membrane potential
TMRE/mitoYFP
****
****
****
4576
4191
100
CCCP
pLenti-Pgs1
WT
Pgs1Crispr
g
Stress-induced Opa1 processing
Untreated
CCCP 5uM 18h
CCCP 20uM 30min
kDa
WT
Pgs1Crisp
WT
Pgs1Crispr
WT
Pgs1Crispr
100
70
55
L-Opa1
S-Opa1
55
Tubulin
h
WT
Opa1Crispr
Opa1CrisprPgs1Crisrpr
Pgs1Crispr
+pLenti-Opa1
+pLenti-Pgs1
+pLenti-Pgs1
Hyperfusion (ActD)
0h
240min
i
Hypertubulation (% cells)
Time (min)
WT
Opa1Crispr
Opa1Crispr+ pLenti Opa1
Opa1CrisprPgs1Crispr
Opa1CrisprPgs1Crispr+ pLenti Pgs1
Pgs1Crispr
Pgs1Crispr + pLenti Pgs1

figure EV5

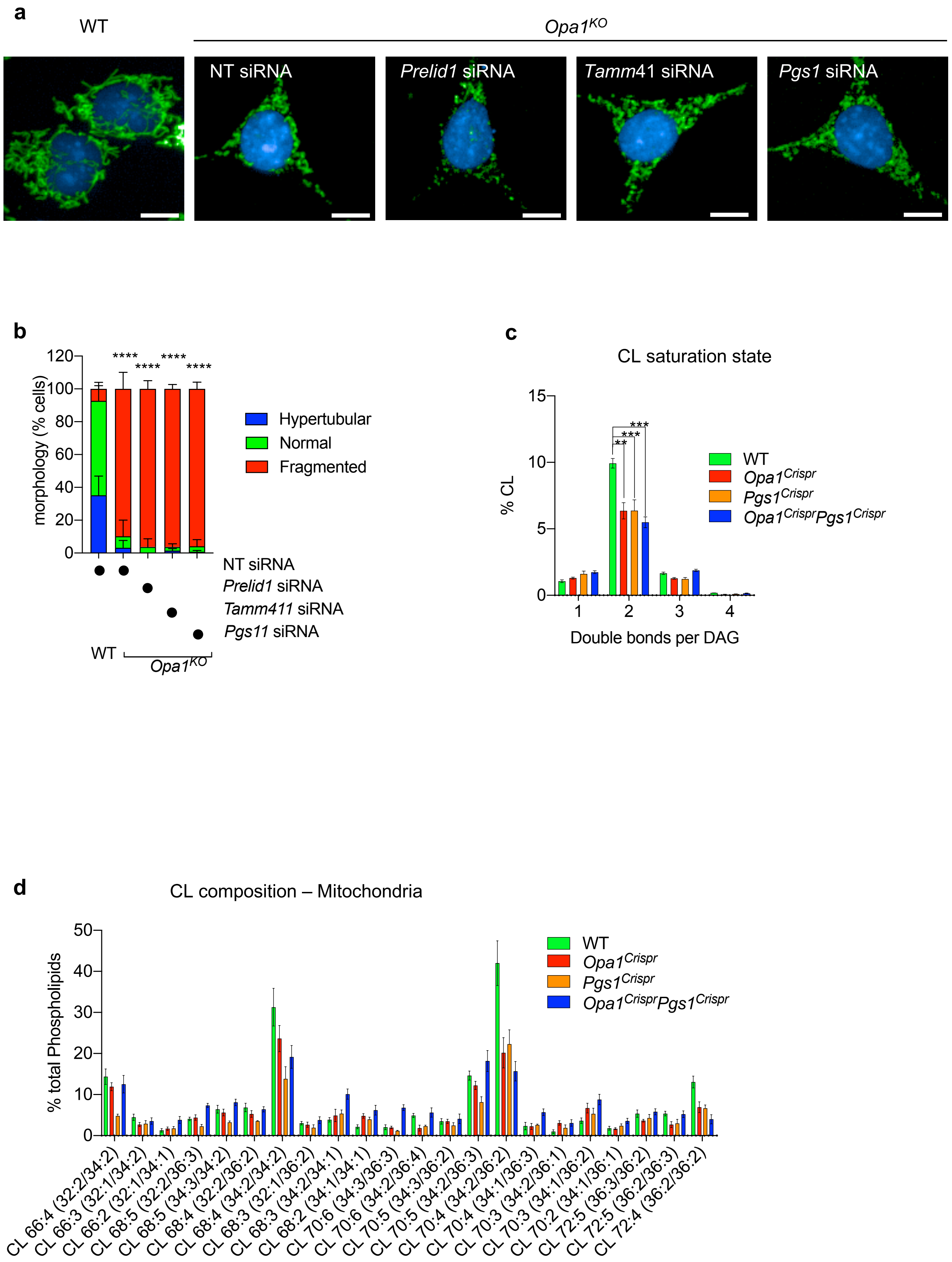
a
WT
Opa1KO
NT siRNA
Prelid1 siRNA
Tamm41 siRNA
Pgs1 siRNA
b
morphology (% cells)
120
100
80
60
40
20
0
****
****
****
****
Hypertubular
Normal
Fragmented
NT siRNA
Prelid1 siRNA
Tamm411 siRNA
Pgs11 siRNA
WT
Opa1KO
c
CL saturation state
% CL
15
10
5
0
**
***
***
WT
Opa1Crispr
Pgs1Crispr
Opa1CrisprPgs1Crispr
1
2
3
4
Double bonds per DAG
d
CL composition – Mitochondria
% total Phospholipids
50
40
30
20
10
0
WT
Opa1Crispr
Pgs1Crispr
Opa1CrisprPgs1Crispr
CL 66:4 (32:2/34:2)
CL 66:3 (32:1/34:2)
CL 66:2 (32:1/34:1)
CL 68:5 (32:2/36:3)
CL 68:5 (34:3/34:2)
CL 68:4 (32:2/36:2)
CL 68:4 (34:2/34:2)
CL 68:3 (32:1/36:2)
CL 68:3 (34:2/34:1)
CL 68:2 (34:1/34:1)
CL 70:6 (34:3/36:3)
CL 70:6 (34:2/36:4)
CL 70:5 (34:3/36:2)
CL 70:5 (34:2/36:3)
CL 70:4 (34:2/36:2)
CL 70:4 (34:1/36:3)
CL 70:3 (34:2/36:1)
CL 70:3 (34:1/36:2)
CL 70:2 (34:1/36:1)
CL 72:5 (36:3/36:2)
CL 72:5 (36:2/36:3)
CL 72:4 (36:2/36:2)

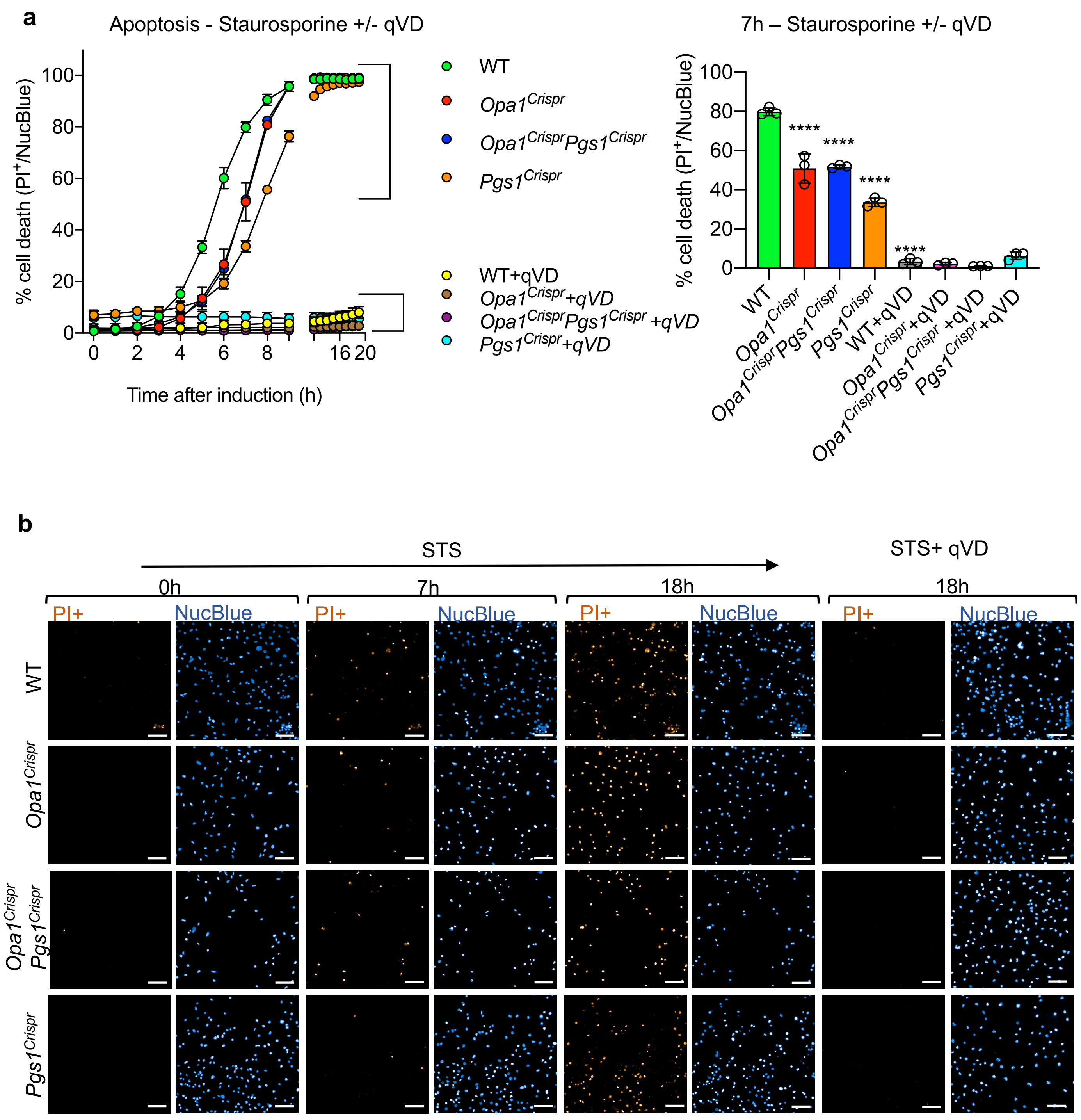
a
Apoptosis - Staurosporine +/- qVD
% cell death (PI+/NucBlue)
Time after induction (h)
WT
Opa1Crispr
Opa1CrisprPgs1Crispr
Pgs1Crispr
WT+qVD
Opa1Crispr+qVD
Opa1CrisprPgs1Crispr +qVD
Pgs1Crispr+qVD
7h – Staurosporine +/- qVD
****
b
STS
STS+ qVD
0h
7h
18h
PI+
NucBlue
WT
Opa1Crispr
Opa1Crispr Pgs1Crispr
Pgs1Crispr

c

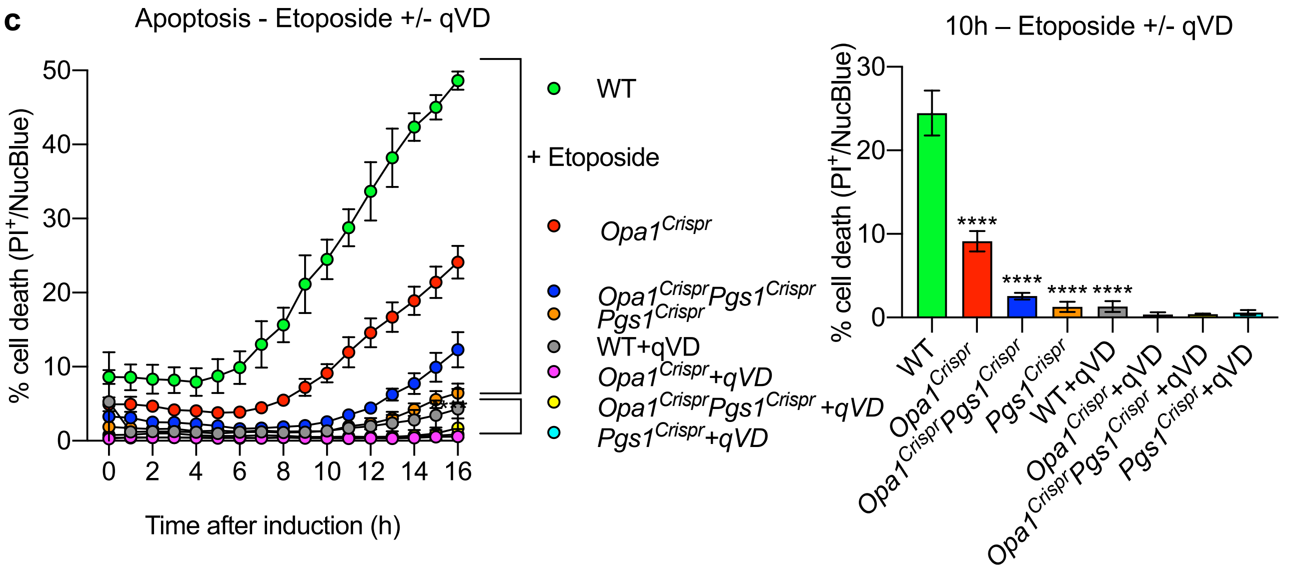
Apoptosis - Etoposide +/- qVD
% cell death (PI+/NucBlue)
Time after induction (h)
0
2
4
6
8
10
12
14
16
0
10
20
30
40
50
WT
+ Etoposide
Opa1Crispr
Opa1CrisprPgs1Crispr
Pgs1Crispr
WT+qVD
Opa1Crispr+qVD
Opa1CrisprPgs1Crispr +qVD
Pgs1Crispr+qVD
10h – Etoposide +/- qVD
****
WT
Opa1Crispr
Opa1CrisprPgs1Crispr
Pgs1Crispr
WT+qVD
Opa1Crispr+qVD
Opa1CrisprPgs1Crispr +qVD
Pgs1Crispr+qVD

d

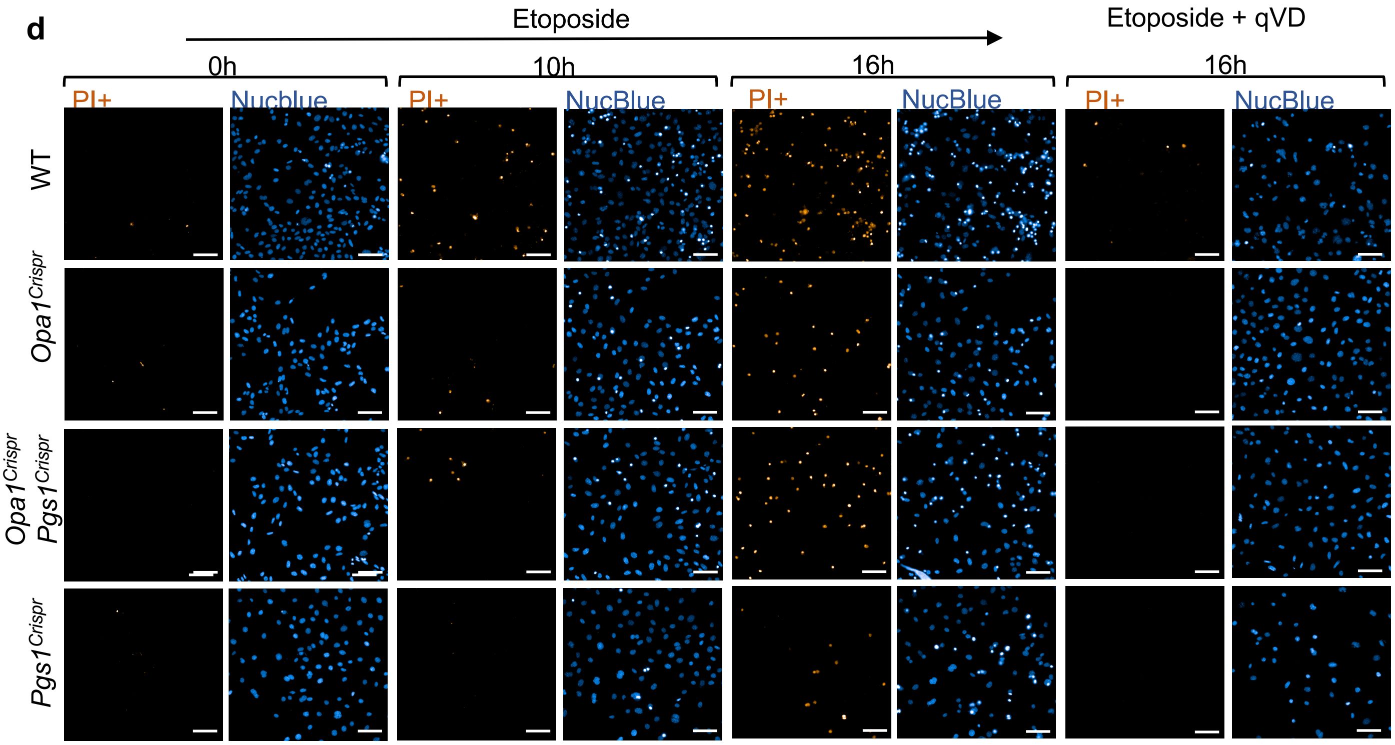
Etoposide
Etoposide + qVD
0h
10h
16h
16h
PI+
Nucblue
NucBlue
WT
Opa1Crispr
Opa1Crispr Pgs1Crispr
Pgs1Crispr

**a** Basal respiration - Oroboros

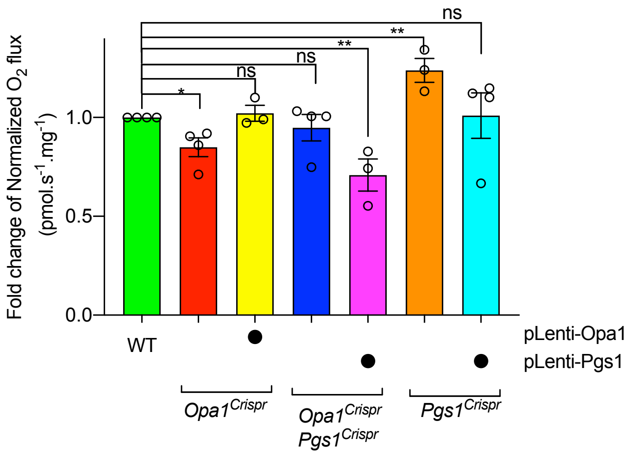


**b** Basal respiration - Seahorse

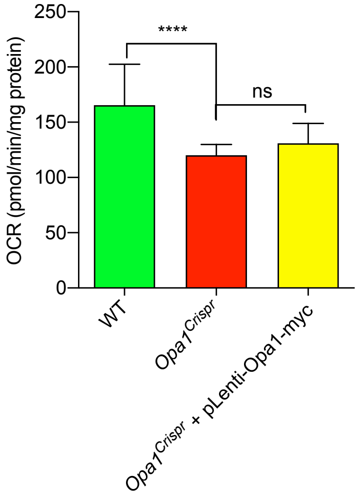


**c** Maximal respiration - Seahorse

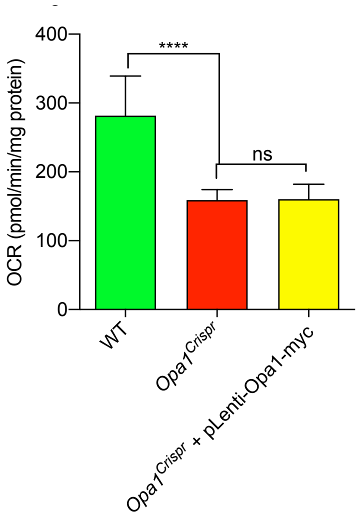
